# Decoding brain activity using a large-scale probabilistic functional-anatomical atlas of human cognition

**DOI:** 10.1101/059618

**Authors:** Timothy N. Rubin, Oluwasanmi Koyejo, Krzysztof J. Gorgolewski, Michael N. Jones, Russell A. Poldrack, Tal Yarkoni

## Abstract

A central goal of cognitive neuroscience is to decode human brain activity--i.e., to infer mental processes from observed patterns of whole-brain activation. Previous decoding efforts have focused on classifying brain activity into a small set of discrete cognitive states. To attain maximal utility, a decoding framework must be open-ended, systematic, and context-sensitive--i.e., capable of interpreting numerous brain states, presented in arbitrary combinations, in light of prior information. Here we take steps towards this objective by introducing a Bayesian decoding framework based on a novel topic model---Generalized Correspondence Latent Dirichlet Allocation---that learns latent topics from a database of over 11,000 published fMRI studies. The model produces highly interpretable, spatially-circumscribed topics that enable flexible decoding of whole-brain images. Importantly, the Bayesian nature of the model allows one to “seed” decoder priors with arbitrary images and text--enabling researchers, for the first time, to generative quantitative, context-sensitive interpretations of whole-brain patterns of brain activity.

## Introduction

A central goal of cognitive neuroscience is to understand how neural and cognitive function interrelate. An important component of this effort is to be able to *decode* cognitive processes from brain activity, or vice versa. Although researchers have dedicated increasing effort to the challenges of brain decoding (Haxby, Connolly, & Guntupalli, 2014; Haynes & Rees, 2006; Kriegeskorte & Kievit, 2013; Mitchell et al., 2004), the vast majority of brain decoding studies to date have focused on fine-grained analysis of a restricted set of cognitive states or experimental tasks--for example, classifying which word or picture a subject is currently perceiving (Cox & Savoy, 2003; Mitchell et al., 2008), or which of several predefined tasks they are engaged in (R. A. Poldrack, Halchenko, & Hanson, 2009; Shirer, Ryali, Rykhlevskaia, Menon, & Greicius, 2012). Such work is notable for its ability to achieve high classification rates of very specific stimuli. However, this accuracy is typically purchased at the cost of high context-specificity: thus far, there is little evidence that the patterns learned by classifiers in such studies can capably generalize to new research sites, experimental designs, and subject populations.

By contrast, much less work has focused on the development of open-ended decoding approaches that are broadly applicable across a variety of contexts. One approach to this type of generalizable decoding is to use large-scale meta-analytic databases such as Neurosynth (Yarkoni et al., 2011) and BrainMap (Laird et al., 2011; Laird, Lancaster, & Fox, 2005) to derive estimates of what a broad variety of brain activations imply about cognitive processing-a form of analysis widely known as *reverse inference* (R. A. Poldrack, 2006; Russell A. Poldrack, 2011). Such efforts necessarily trade fidelity for breadth; that is, they allow researchers to draw inferences about almost any cognitive process that has been frequently studied with fMRI, but these inference are coarse, and come with a high degree of uncertainty. An illustrative study was conducted by Chang et al. (2012), who used the Neurosynth database to "decode" the functional correlates of three distinct right insula clusters. The analytical strategy involved correlating each insula map with dozens of Neurosynth meta-analysis maps and drawing conclusions about function based on differences in relative similarity (e.g., an anterior insula region showed greatest similarity to executive control-related meta-analysis maps; a ventral insula region showed greatest similarity to affect-related maps; etc.). Other studies have used a similar approach to infer the putative functional correlates of whole-brain maps in a variety of other settings (e.g., Andrews-Hanna, Saxe, & Yarkoni, 2014; Bzdok et al., 2013; Smith et al., 2009).

More recently, we have generalized this approach and implemented it in the online Neurosynth and NeuroVault platforms (http://neurosynth.org/decode). At present, researchers can upload arbitrary whole-brain maps to the NeuroVault repository and instantly decode them against the entire Neurosynth database. This decoding functionality provides researchers with a quantitative means of interpreting whole-brain activity patterns--potentially replacing the qualitative conclusions more commonly drawn in the literature. However, the present approach--which is based entirely on computation of spatial similarity coefficients between the input map and comparison meta-analysis maps--has several weaknesses that limit its utility as a general-purpose decoding framework. Chief among these is that the approach is not grounded in a formal model: it allows one to estimate the similarity of any given brain activity map to other canonical maps, but does not provide a principled way to interpret these mappings.

Furthermore, the approach does not attempt to identify any latent structure that presumably makes such mappings useful--for example, individual brain regions or functional brain networks that correspond to specific cognitive processes. A generative framework for decoding brain activity would offer researchers a number of important benefits: it would facilitate the learning of interpretable latent structures from a mass of superficial brain-cognition mappings; provide the ability to decode bidirectionally--i.e., to not only identify functional correlates of arbitrary whole-brain images, but to also project descriptions of experimental tasks or psychological concepts into image space; and allow principled generation of novel exemplars or combinations of events that have never been seen before (e.g., what pattern of brain activity would a task combining painful stimulation and phonological awareness produce?).

Perhaps most importantly, by virtue of explicitly modeling both the joint and marginal probabilities of all events, a generative framework would provide the ability to contextualize predictions through the explicit use of Bayesian priors. At present, all brain decoding approaches we are aware of are acontextual: they provide researchers with no way to integrate contextual information or prior belief into the decoding process. Since many if not most brain regions are generally understood to contain multiple circuits with potentially distinguishable functions, knowledge of the experimental context within which a pattern of brain activity unfolds should, in principle, constrain interpretation of observed brain activity. Left inferior frontal gyrus activation may mean different things in the context of language comprehension (Vigneau et al., 2006), emotion regulation (Buhle et al., 2014), or response inhibition (Swick, Ashley, & Turken, 2011). More generally, true reverse inference--i.e., the move to draw conclusions about the likelihood of different mental states conditional on observed brain activity--is an inherently Bayesian notion that requires one to formally model (and specify) the prior probability of each term or concept’s occurrence. Whereas a similarity-based decoding approach cannot easily support such specification, it is intrinsic to a generative model.

Here we take the first steps towards these goals by introducing a generative Bayesian decoding framework based on a novel topic model---Generalized Correspondence Latent Dirichlet Allocation (GC-LDA)---that learns latent topics from the meta-analytic Neurosynth database of over 11,000 published fMRI studies (Yarkoni et al., 2011). GC-LDA generates topics that are simultaneously constrained by both anatomical and functional considerations: each topic defines a spatial region in the brain that is associated with a highly interpretable, coherent set of cognitive terms. We demonstrate that the dictionary of topics produced by the GC-LDA model successfully captures known anatomical and functional distinctions and provides a novel data-driven metric of hemispheric specialization. We then take advantage of the topic model’s joint spatial and semantic constraints to develop a bidirectional, open-ended decoding framework. That is, we demonstrate the ability to extract both a text-based representation of any whole-brain image, and a whole-brain activity pattern corresponding to arbitrary text. Importantly, the Bayesian nature of the model allows us to formally specify a decoder’s priors by "seeding" it with any arbitrary combination of images and text. The direct consequence is that, for the first time, researchers are able to generative quantitative, context-sensitive interpretations of whole-brain patterns of brain activity.

## Results

### Mapping the functional neuroanatomy of the brain with topic models

Our decoding framework is built on a widely-used Bayesian modeling approach known as *topic modeling* (Blei, 2012; Blei, Ng, & Jordan, 2003). Topic modeling is a dimensionality-reduction technique, which decomposes a corpus of documents into a set of semantically coherent probability distributions over words, known as *topics*. Given this set of topics, each document can be represented as a probabilistic mixture of topics. Topic models have been successfully applied to a wide range of problems, including text classification <u>(Mcauliffe and Blei 2008: Rubin et al. 2011)</u> information retrieval (Zhai, Chengxiang, & John, 2001), image classification (Cao, Liangliang, & Li, 2007), and theme discovery (Griffiths & Steyvers, 2004; Steyvers, Smyth, Rosen-Zvi, & Griffiths, 2004), and are now regarded as a standard technique for text and image analysis. An important feature from a decoding standpoint is that topic models are generative in nature: they allow a principled approach for bidirectional mapping from documents to latent components and vice versa; probabilistic generation of entirely new (i.e., previously unseen) documents; and formal Bayesian updating that can allow for explicit specification of the prior topic probabilities. We return to these features later.

**Figure 1.**
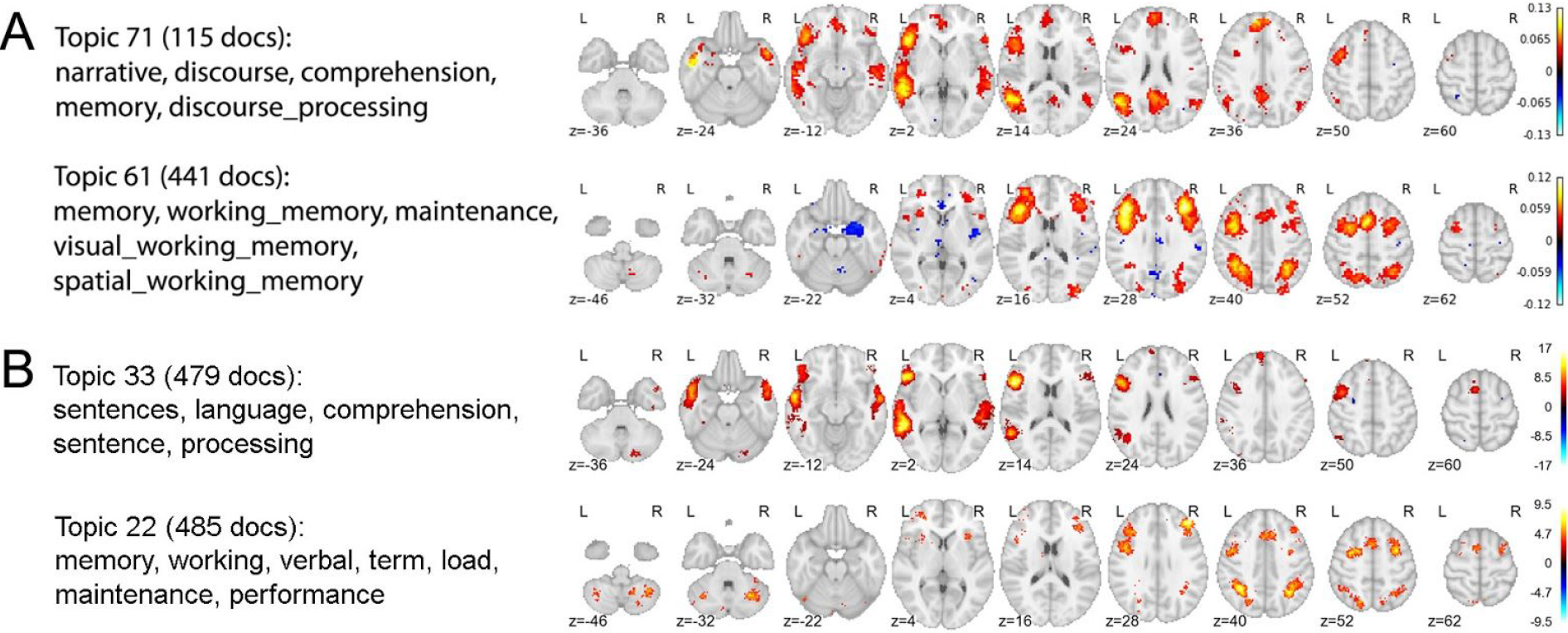
Replication of topics from Poldrack et al. (2012). Figure shows the results of applying the generic LDA model (Blei et al., 2003) to the Neurosynth database, as described in Poldrack et al. (2012). (A) Selected topics reported in Poldrack et al. (2012) using an older Neurosynth database of 5,809 studies. (B) Closest matching topics when applying the same approach to the current, expanded, Neurosynth database (11,406 studies).

In previous work, we used a standard topic model to extract 200 semantically coherent topics from the abstracts of all published fMRI articles contained in an older and smaller version of the Neurosynth database (5,809 studies; Russell A. Poldrack et al., 2012). We then projected each topic onto the space of brain activity to identify brain regions associated with distinct cognitive profiles. A direct replication of this earlier approach using the current, and much larger, Neurosynth database (11,406 studies) produces very similar results (e.g., Figure 1). As Figure 1 illustrates, the structure-function mappings produced by this approach converge closely with numerous other findings in the literature---e.g., the presence of a strongly left-lateralized language network (Vigneau et al., 2006) and the involvement of dorsal frontoparietal regions in working memory and executive control (Duncan, 2010). However, because the standard topic model operates only on the text of publications, the topics it produces are not constrained in any way by neural data. Furthermore, the spatial mappings for each topic are indirectly computed via the documents’ topic loadings--the spatial data is not built into the model. The result is a set of widely distributed, network-like activation maps that closely resemble the whole-brain maps produced by individual fMRI experiments. While such an approach is informative if one’s goal is to identify the distributed neural correlates of coherent psychological topics, it is of little help in the search for relatively simple, well-defined functional-anatomical atoms. A similar limitation applies to more recent work by Yeo et al, who used a more sophisticated topic model to derive a set of *cognitive components* that map in a many-to-many fashion onto both behavioral tasks and patterns of brain activity (Yeo et al., 2014). While the latter approach represents an important advance in its simultaneous use of both behavioral and brain activity data, the resulting spatial components remain relatively widely distributed, and do not provide insight into the likely cognitive roles of well-localized brain regions.

### The GC-LDA model

To extract structure-to-function mappings focused on a more granular, region-like level of analysis, we developed a novel topic model based on the Correspondence-LDA model (Blei & Jordan, 2003) that generates topics simultaneously constrained by both semantic and spatial information. We term this the Generalized Correspondence LDA (GC-LDA) model <u>(Fig. 2: for details, see Rubin. T. N., Koveio. O., Jones. M. N…)</u>. The GC-LDA model learns a set of latent topics, each associated with (i) a spatial probability distribution over brain activations and (ii) a set of words that co-occur in article abstracts. This extension of the Correspondence-LDA model incorporates a flexible spatial distribution that can be adjusted according to the goals of the experimenter. Here we focus on a version of the model in which each topic is associated with a mixture of two Gaussian clusters constrained to show symmetry around the x-axis, enabling us to directly quantify the degree of hemispheric symmetry (or lack thereof) displayed by each topic.

**Figure 2.**
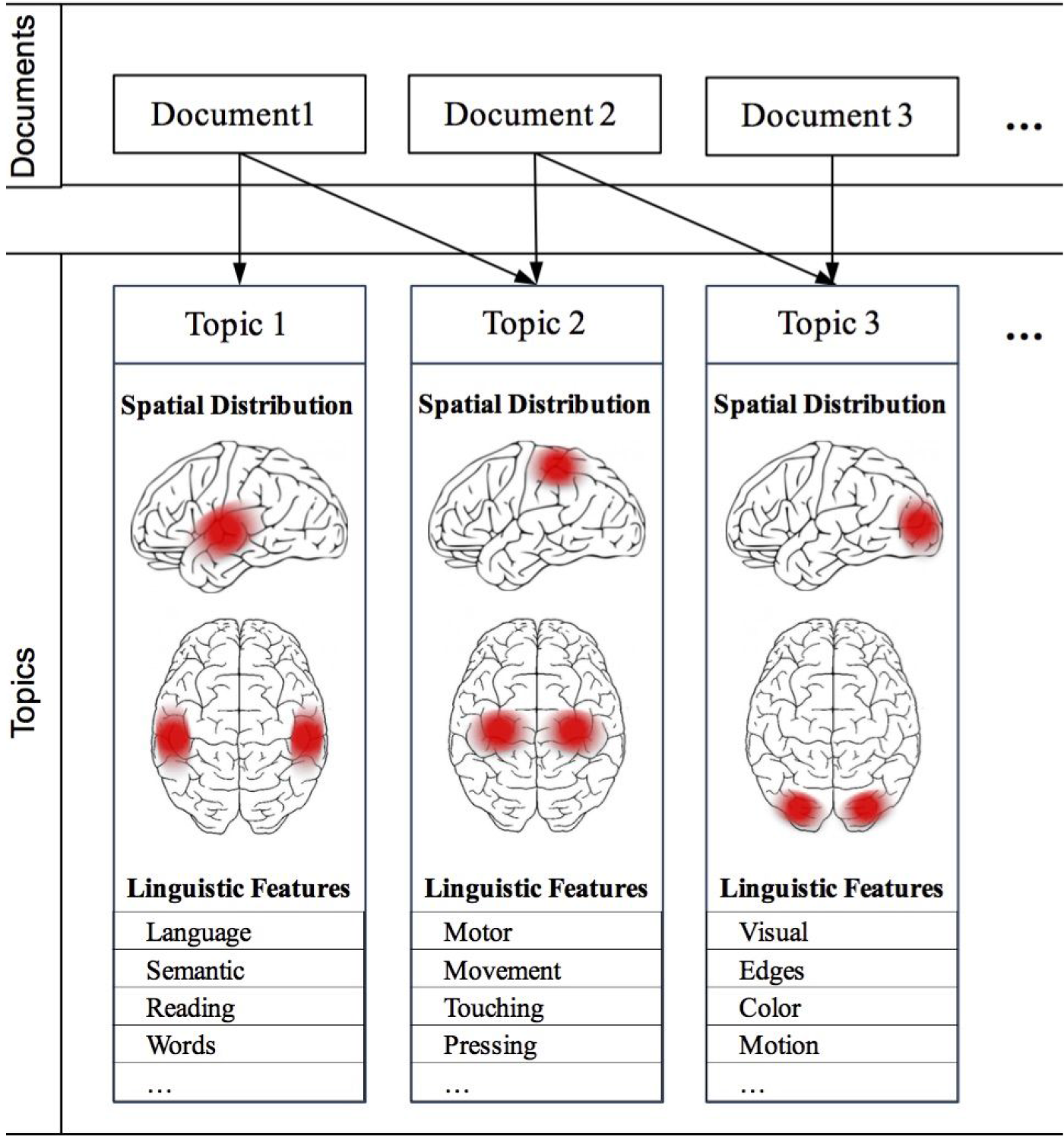
Schematic overview of the GC-LDA model. Each document (an article in the Neurosynth corpus) is represented as a mixture of learned latent topics, where each topic is associated with both a 3-dimensional Gaussian spatial distribution, and a set of linguistic terms extracted from the abstract text.

Figure 3 displays selected topics extracted using the GC-LDA model (for comprehensive results, see Supplementary Figure 1 and neurovault.org/collections/EBAYVDBZ/). As illustrated, the model produced numerous topics that had well-defined joint spatial and semantic representations (Fig. 3a)--approximately half of the 200 extracted topics were clearly interpretable (see Supp. Fig. 1 for full details). Many of these topics successfully captured relatively basic associations between specific structures and their putative functions; for example, we identified topics associated with amygdala activation and emotion; reward and the ventral striatum; hippocampus and memory; fusiform face area and face perception; and motion perception and the V5/MT complex, among others (Figure 3b). In other cases, the model successfully captured and localized higher-level cognitive processes--e.g., topics associated with the temporoparietal junction and mentalizing, temporal pole and person perception, or ventromedial PFC and valuation, among others (Fig. 3b). In supplementary analyses, we further demonstrate that the full set of 200 topics can be used to accurately “reconstruct” arbitrary patterns of whole-brain activity, providing an interpretable, low-dimensional way to summarize virtually any whole-brain image (Supplementary Results).

**Figure 3.**
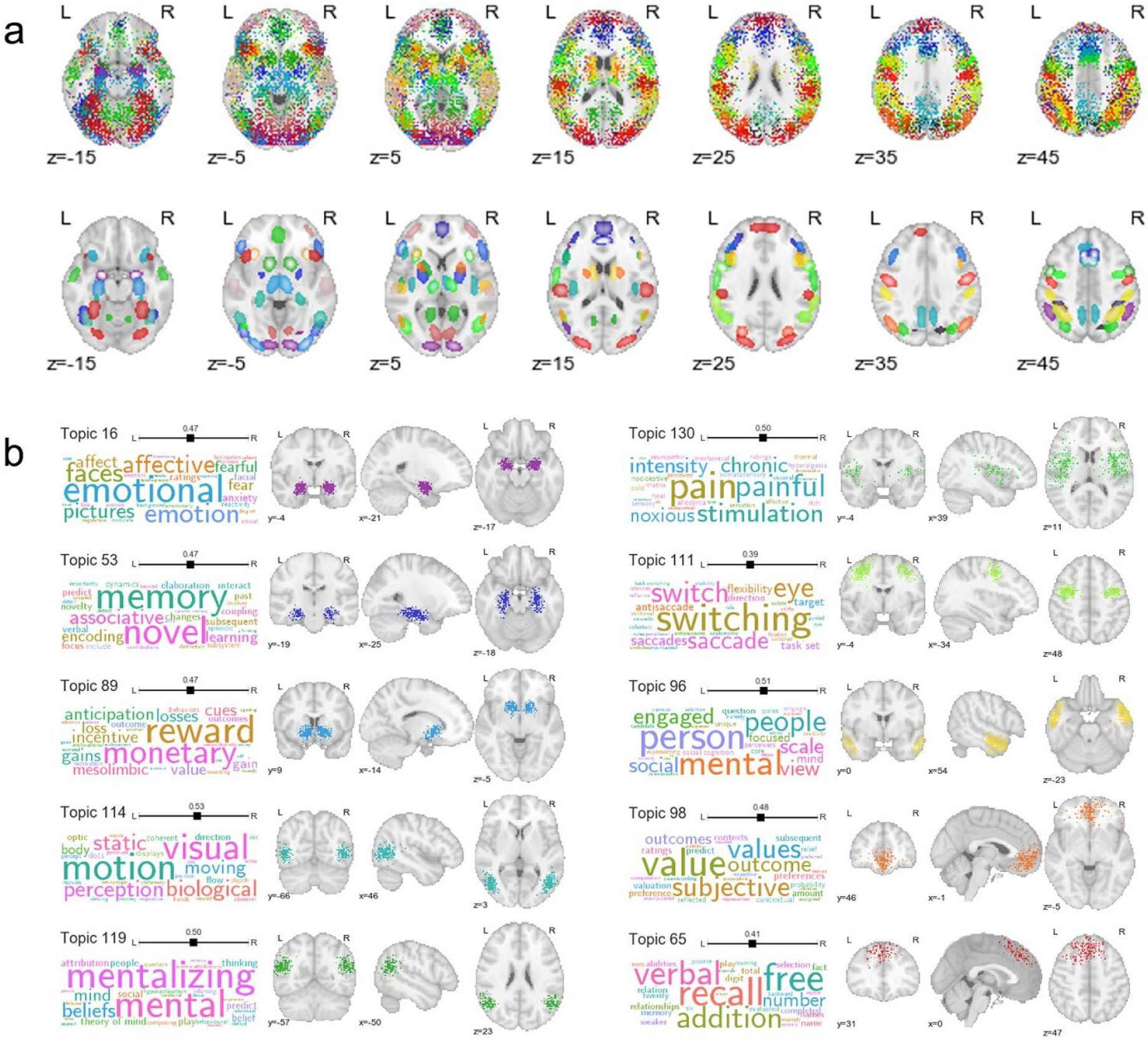
Selected topics learned by the GC-LDA model (for full results, see Supplementary Figure 1). (a) Spatial distributions for 90 of the 200 topics. Each color represents a different topic. Top row: hard assignments of activations to topics; each point represents a single activation from a single study in the Neurosynth database. Bottom row: estimated multivariate Gaussian mixture distribution of each topic, (b) Top semantic associates (word clouds) and activation distributions (brain orthviews) for selected topics. The size of a term in each word cloud is proportional to the strength of loading on the corresponding topic.

### Probabilistic structure-to-function mapping

An important feature of the GC-LDA model is that it is probabilistic, and avoids the common, but restrictive, clustering assumption that each voxel should only be assigned to a single group (Bellec, Rosa-Neto, Lyttelton, Benali, & Evans, 2010; Blumensath et al., 2013; Craddock, James, Holtzheimer, Hu, & Mayberg, 2012; Power et al., 2011; Yeo et al., 2011). By allowing extracted topics to overlap with one another in space, the model explicitly acknowledges that the brain contains spatially overlapping circuits with thematically related functions. Figure 4 illustrates the close spatial and semantic relationships between 10 different topics localized to overlapping parts of the parietal cortex along the banks of the intraparietal sulcus (IPS). Note the particularly similar posterior parietal cortex (PPC) distributions of topics associated with visuospatial processing, working memory, and general task engagement. These results are consistent with electrophysiological findings of highly heterogeneous, and typically complex, response profiles in PPC neurons (including coding of visual object location, direction of attention, motor plans, etc.; Andersen, Essick, & Siegel, 1985; Bushnell, Goldberg, & Robinson, 1981; Mountcastle, Lynch, Georgopoulos, Sakata, & Acuna, 1975; Snyder, Batista, & Andersen, 1997), and underscore the difficulty individual fMRI studies may face in trying to isolate brain-cognition mappings via a hemodynamic signal that sums over millions of neurons at each voxel.

**Figure 4.**
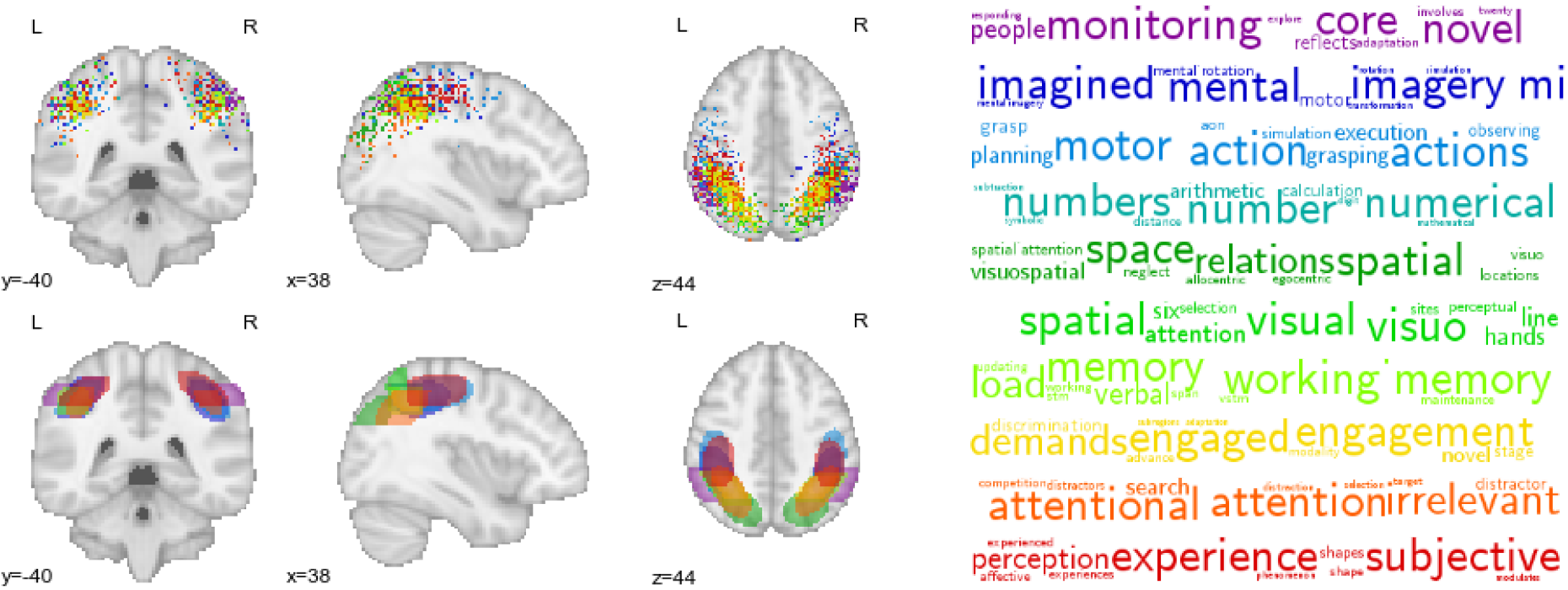
Activation profiles and top-loading words for spatially overlapping topics in parietal cortex. Top row: hard assignments of activations to topics; each point represents a single activation from a single study in the Neurosynth database. Bottom row: estimated multivariate Gaussian mixture distribution of each topic.

Analogously, the probabilistic nature of the GC-LDA mappings can also provide insights into the compositional character of most cognitive states---i.e., the fact that most states are likely to recruit activation of a number of spatially distinct brain regions. Figure 5 displays activation and word distributions for a number of emotion-related topics. Different topics captured different aspects of emotional processing: consistent with extensive previous work, extrastriate visual cortex and amygdala were associated with perceptual processing of emotion (LeDoux, 2003; Phelps, 2006; Zald, 2003); rostral anterior cingulate cortex and anterior insula were associated with experiential aspects of emotion (Lindquist, Wager, Kober, Bliss-Moreau, & Barrett, 2012); and lateral frontal cortex was associated with emotion regulation (Buhle et al., 2014; Kohn et al., 2014).

**Figure 5.**
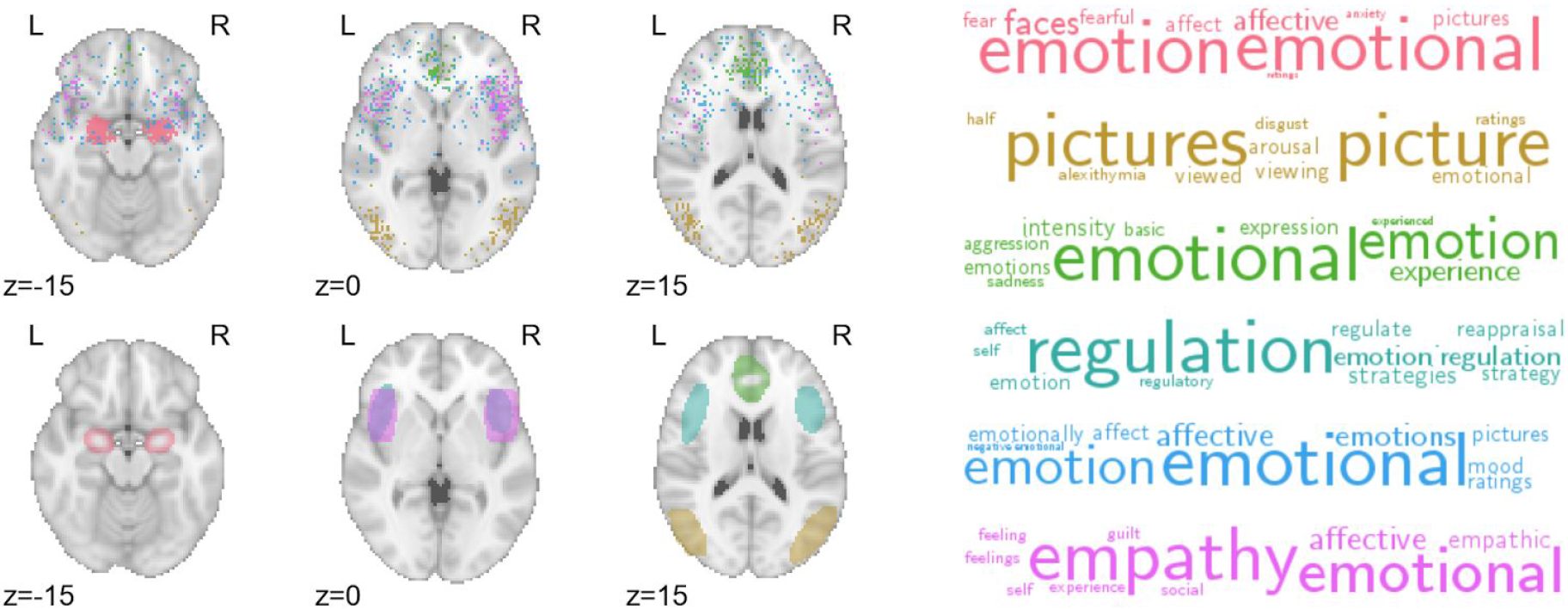
Activation profiles and toploading words for emotionrelated topics. Top row: hard assignments of activations to topics; each point represents a single activation from a single study in the Neurosynth database. Bottom row: estimated multivariate Gaussian mixture distribution of each topic.

### A data-driven window into lateralization of function

As noted above, each topic in the GC-LDA model was deliberately constrained to reflect two sub-regions reflected around the brain’s x-axis. This constraint allowed us to estimate the relative weight of activations for each topic in the left vs. right hemisphere--in effect providing a novel, data-driven index of hemispheric specialization. As one might expect given the marked degree of activation symmetry observed in most fMRI studies, most topics showed little or no hemispheric bias (Figure 6, top). However, there were a number of notable exceptions (e.g., Fig. 6, bottom). Several language-related topics localized strongly to left-hemisphere language regions—including inferior and middle frontal gyrus, posterior superior temporal sulcus, and inferotemporal cortex (encompassing the putative visual word form area; Dehaene, Stanislas, & Laurent, 2011). Right-lateralized topics were fewer in number and generally showed a weaker hemispheric asymmetry, but notably included a face processing topic localized to the putative fusiform face area (Kanwisher, McDermott, & Chun, 1997), and an inhibitory control-related topic localized to the right ventral anterior insula (Aron, Robbins, & Poldrack, 2004). To our knowledge, these findings constitute the first data-driven estimation of region-level functional hemispheric asymmetry across the whole brain.

**Figure 6.**
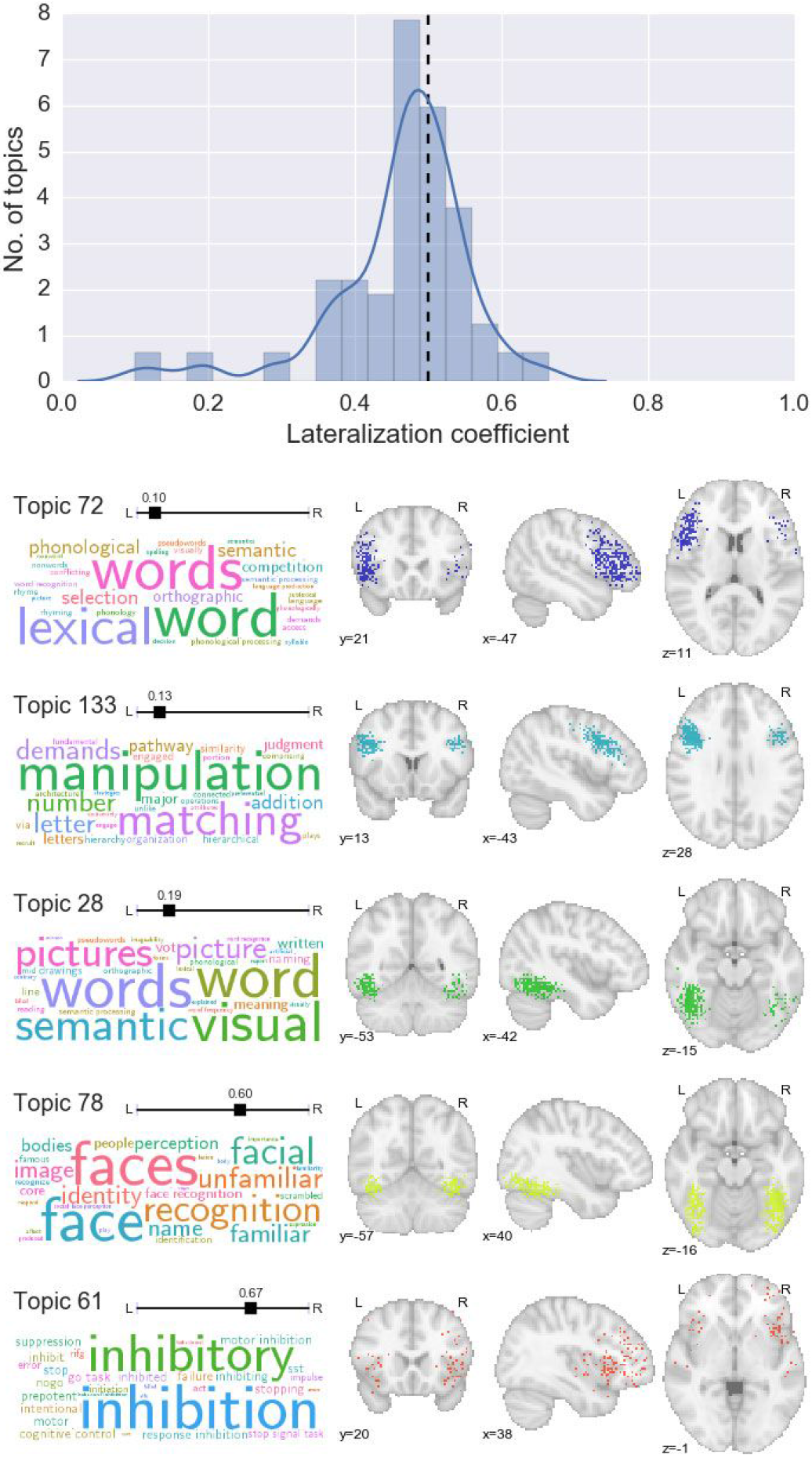
Data-driven estimation of hemispheric lateralization of cognitive function. Top: histogram and kernel density estimation plot of the lateralization coefficient for all topics. Values below 0.5 represent left-lateralization; values above 0.5 represent right-lateralization. Bottom: selected topics that displayed notable hemispheric lateralization.

### Automatic text-to-image and image-to-text decoding

Importantly the GC-LDA model is able to produce probabilistic estimates of word and activation distributions for entirely new data points. Moreover, because each topic is associated with both a word distribution and a spatial distribution, we can proceed bidirectionally---either translating arbitrary text into image space, or decoding activations or images for their associated semantic content. Figure 7 illustrates three different applications of this approach. First, we can generate estimated activation probabilities for any word or set of words. Figure 7A illustrates three concrete examples. In (1), we observe a complex, distributed pattern of activity for the term ’motor’, including activations in primary and supplementary motor cortices, cerebellum, and the basal ganglia. This result demonstrates that even though each topic in our dictionary is spatially constrained, individual words will often still have widely distributed neural correlates by virtue of loading on multiple topics.

In (2) we pass in a list of generic cognitive effort-related terms (’effort’, ’difficult’, and ’demands’), and observe highly circumscribed activations in frontoparietal regions frequently implicated in general goal-directed processing (Dosenbach et al., 2006; Duncan, 2010). This result demonstrates the GC-LDA model’s ability to produce topics with relatively abstract semantics: while few studies explicitly set out to study the neural correlates of task difficulty or cognitive effort, our model successfully learns that regions like anterior insula and preSMA--which tend to activate in a very wide range of studies--likely support fairly general cognitive operations non-selectively invoked by many different tasks (cf. Chang et al., 2012; Nelson et al., 2010; Neurosynth; Yarkoni et al., 2011).

Lastly, in (3), we provide a full sentence as input ("painful stimulation during a language task"), producing a map with peaks in both pain-related (e.g., posterior insula) and language-related left perisylvian regions. While the model follows the bag-of-words assumption (i.e., the order of words has no effect on the generated image), its compositional character is evident, in that it is possible to generate a predicted image for virtually any cognitive state or states that can be described in text.

Second, we can generate a list of plausible semantic associates for any set of discrete brain coordinates. Figure 7B illustrates how this approach can be used to probe the function of a particular region both in isolation and in context. The top row lists the top probabilistic word associates for a temporoparietal region centered on MNI x-y-z coordinates (−56, −52, 18). The function of this region appears ambiguous---likely reflecting the presence of multiple overlapping neural circuits---with the top associates including ’reading’, ’mentalizing’, and ’pictures’. However, adding other coordinates strongly constrains functional interpretation. The addition of medial parietal (0, −58, 38) and dorsomedial prefrontal (4, 54, 26) activations (middle row) produces strong overall loadings on default network-related terms such as ’self’, ’social’, and ’moral’. By contrast, adding left superior temporal sulcus (−54, −40, 0) and left inferior frontal gyrus (−50, 26, 6) activations (bottom row) instead produces strong loadings on language and reading-related terms (Fig. 7B). Thus, the GC-LDA model allows researchers to freely explore structure-function mappings in the brain in a context-specific way that recognizes that the cognitive operations supported by individual regions can contribute to multiple distinct cognitive functions.

Lastly, and perhaps most powerfully, the activation-to-word mapping approach can be generalized to entire whole-brain images. Given any real-valued input image, we can use the GC-LDA topics to generate a rank-ordered list of associated terms. While the output values cannot be interpreted as actual probabilities (due to the arbitrary scale of the inputs), the results are highly informative, providing a quantitative, literature-based decoding of virtually any pattern of whole-brain activity. Figure 7C illustrates the results for selected images, including two of the cognitive components from Yeo et al. (2014), two of the BrainMap ICA components from Smith et al. (2009), and two group-level HCP task contrasts (for additional results, see Supplementary Figures 2 and 3). The decoded term list converges closely with extensive prior work; for example, a BrainMap ICA component focused largely on extrastriate visual cortex and adjacent inferotemporal areas is associated with motion, face perception, and other vision-related terms; a cognitive component from Yeo et al. (2014) largely co-extensive with the frontoparietal control network loads most strongly on terms like “working memory”, “demands”, and “numerical”, and so on.

**Figure 7.**
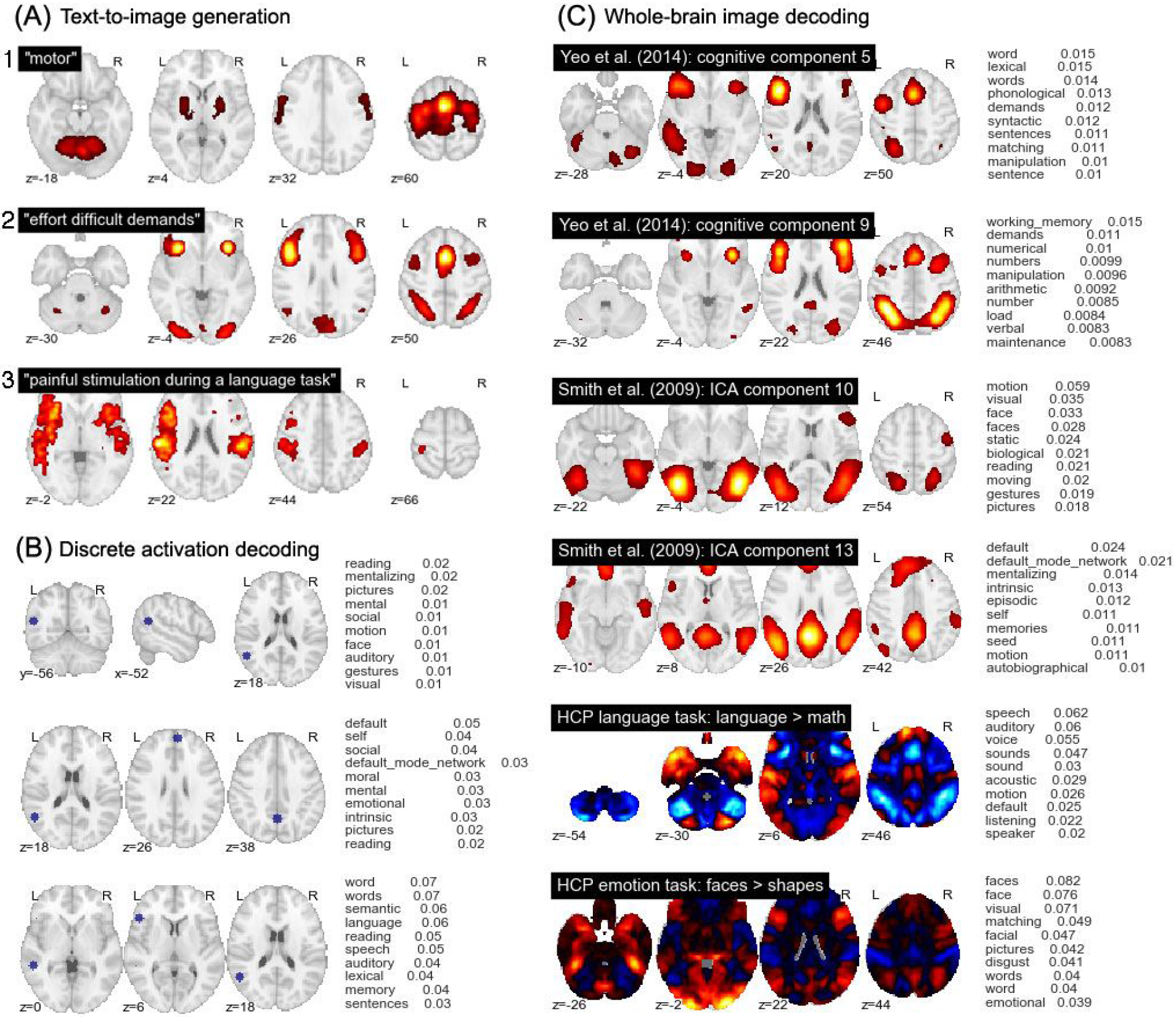
Examples of generative text-to-image and image-to-text mapping using the trained GC-LDA model. (A) Generation of predicted whole-brain images from arbitrary text. (B) Topic-based decoding of discrete activation coordinates. (C) Topic-based decoding of continuous whole-brain images; examples selected from the cognitive components reported in Yeo et al. (2014), the BrainMap ICA components reported in Smith et al. (2009), and the language and emotion contrasts from the n = 500 release of the HCP dataset. Note that the scale of the values in (B) and (C) is dependent on the input image, and should not be assigned an absolute interpretation.

To more formally assess the performance of the decoder in an unbiased way, we used a set of NeuroVault images that were previously manually annotated using labels derived from the Cognitive Atlas ontology (Russell A. Poldrack et al., 2011). For each image, we used the image-to-text decoder to generate an image-specific rank-ordering of the 1,000 most common terms in the entire Neurosynth corpus. We then identified the rank, within that list, of each human-annotated Cognitive Atlas label. The median rank across all 300 images was 220--an impressive value considering the open-ended nature of the task and the unfiltered nature of the NeuroVault database (i.e., there is no guarantee that the images uploaded to Neurovault actually reflect the processes they are intended to reflect--a point we discuss further in the next section). By comparison, when we generated a null distribution of 1,000 permutations and computed the same median statistic, the mean and minimum values across all permutations were 442 and 384, respectively. In other words, the decoder produced rankings that were vastly more similar to expert human judgments than one would expect by chance.

### Brain decoding in context

Importantly, the above analysis provides a necessarily conservative estimate of the performance of our decoder, because in many cases, the discrepancy between human-annotated and automatically-decoded labels is bound to reflect error in the former rather than the latter. We note that human-generated annotations typically reflect researchers’ beliefs about which cognitive processes a particular experimental manipulation is *supposed* to influence, and do not represent ground truth. For example, the HCP Gambling Task (adapted from Delgado, Nystrom, Fissell, Noll, & Fiez, 2000) was putatively designed "to assess reward processing and decision making" (Barch et al., 2013). Yet the contrast between the reward and loss conditions (depicted in Fig. 8) reveals robust reward-related increases in visual and frontoparietal cortices (Fig. 8, top). Not surprisingly, terms like ’visual’, and ’working memory’ are at the top of the list returned by our decoder (see “uniform prior” results in Fig. 8). Does this mean that the decoder is performing poorly, and failing to recover a known ground truth? No. Given the non-canonical pattern of observed brain activity, we believe a more plausible alternative is that the manipulation in question simply had a more complex effect on cognition than the "Reward vs. Loss" label might lead one to expect. In other words, the "assumption of pure insertion"--i.e., that the gain vs. loss contrast measures only cognitive processes related to reward or loss processing--is probably unwarranted in this case, as in many others (Friston et al., 1996; Russell A. Poldrack, 2010).

The potential for discrepancy between expert human judgment and automated decoding creates an interesting conundrum: which answer should a neuroimaging researcher trust? Our view is that there is no blanket answer to this question; much depends on the particular context. Importantly, our decoding framework provides a way to quantitatively synthesize researchers’ prior beliefs with the associations learned by the GC-LDA topic model by explicitly manipulating the prior probabilities of the 200 topics. Because our model allows for bi-directional decoding (text-to-image or image-to-text), topic priors can be set by “seeding” the model with either a whole-brain image (or images), or a set of terms. The seeds are decoded in the normal way to update the initial uniform prior, and subsequent decoding is then based on the updated (non-uniform) priors. The approach is illustrated in Figure 8, which displays the results of a topic decoding analysis for two HCP task contrasts when the decoder is seeded (i) with uniform priors, (ii) with a set of reward-related terms, or (iii) with the whole-brain Neurosynth meta-analysis map for the term “reward” (http://neurosynth.org/analyses/terms/reward). The strength of the prior is also explicitly varied.

The major result illustrated in Fig. 8 is that if one is able to specify a prior belief about the experimental context, the decoder respects this prior and produces results that are, to varying degrees, biased in the direction of the prior. The decoder results are implicitly smoothed by the underlying latent topics; for instance, in the top row of Fig. 8, the terms “monetary” and “anticipation” appear near the top of the text-seeded results, even though they were not included in the list of seed terms. Moreover, the priors do not overwhelm the data (unless the strength parameter is set very high, as in the columns with weight = 0.25). When the reward-related priors are applied to a map that is highly inconsistent with the prior--as in the Language > Math contrast in the bottom row of Fig. 8--the change in decoder results is much more subtle. Thus, our decoding framework provides a quantitative way of contextualizing interpretations of fMRI data in a principled way--or, alternatively, assessing the degree to which a particular interpretation is dependent on typically unstated prior beliefs.

**Figure 8.**
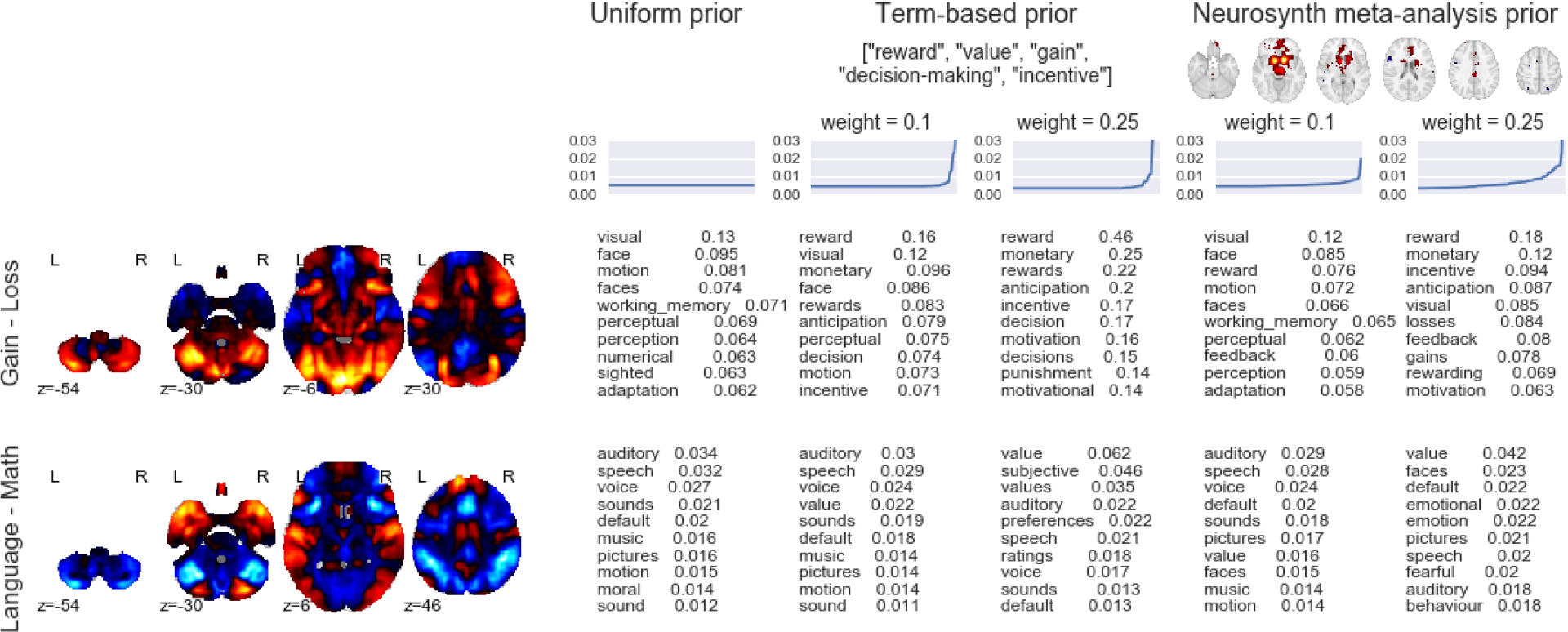
Effects of different topic priors on decoding results. The top 10 terms produced by the decoder are displayed for two different HCP contrasts (Gain > Loss from the Gambling task and Language > Math from the Language task) and three different sets of topic priors (left: uniform prior; middle: priors seeded with a list of reward-related terms; right: priors seeded with the Neurosynth “reward” meta-analysis map). For the non-uniform priors, results are displayed for priors of differing strengths (weak = 0.1, strong = 0.25). Line plots above the decoder outputs illustrate the prior distribution of topics used in each analysis (for the sake of visual clarity, topics are ordered by increasing weight separately in each case).

## Discussion

The present work significantly advances beyond previous efforts with respect to both (a) the modeling of the latent structure of neurocognition and (b) the open-ended decoding of human brain activity. With respect to the former, the GC-LDA topic model we developed introduces several innovative features to the literature. First, the simultaneous use of spatial and semantic information allows the model to learn topics that have both well-localized spatial representations, and clear semantic correlates. Approximately half of the 200 topics we extracted in a completely data-driven way closely tracked previous functional and anatomical distinctions reported in previous fMRI studies. Second, the probabilistic nature of the resulting topics stands in contrast to many previous clustering and parcellation approaches, and more accurately reflects the many-to-many nature of the relationship between cognitive constructs and neurobiological structures. Third, the GC-LDA model’s spatial symmetry constraint enabled us to generate brain-wide, data-driven estimates of the relative hemispheric lateralization of distinct cognitive topics. Consistent with the broader literature, most topics displayed a high degree of symmetry, with notable exceptions including the strong left-lateralization of language- and memory-related topics, and the more modest right-lateralization of response inhibition and face-related topics. Finally, the spatially compact, semantically well-defined nature of the 200 extracted topics makes the full topic set an ideal basis set for use in dimensionality reduction and image interpretation applications (as exemplified by the “topic reconstruction” analyses reported in the Supplementary Results and illustrated in Supplementary Figures 4-7).

From the standpoint of efforts to decode whole-brain activation patterns, our results also advances beyond previous work. First, by simultaneously constraining topics both spatially and semantically, the GC-LDA model generates topics designed to maximize the correspondence between cognition and brain activity. By contrast, previous open-ended decoding approaches have typically relied on predefined cognitive ontologies (Bzdok et al., 2013; Smith et al., 2009) or terms (Andrews-Hanna et al., 2014; Chang et al., 2012), which are unlikely to collectively maximize the parsimony of derived brain-cognition mappings. Second, the generative nature of our decoding framework facilitates bidirectional decoding, enabling researchers not only to identify likely functional correlates of whole-brain activity patterns or sets of discrete activations, but also to project flexible text descriptions of tasks or processes into image space.

Third, our Bayesian approach allows researchers to formally specify priors on the GC-LDA topics, providing a powerful means of contextualizing interpretations and accounting for prior expectations and beliefs. We illustrate how a researcher can flexibly "seed" a decoding analysis using cognitive terms and/or whole-brain maps, thus ensuring that the decoder respects prior information about the experimental context. Current decoding approaches are forced to rely on unstated and inflexible assumptions about the base rates associated with different cognitive processes or tasks--a limitation that makes it difficult to know how much trust to place in a particular interpretation of one’s results. While our Bayesian updating approach currently has important limitations (see below), it represents an important step towards the goal of being able to decode arbitrary patterns of whole-brain activity in a way that formally synthesizes prior knowledge with observed results.

Naturally, the present work remains constrained by a number of important limitations. First, the specificity of the extracted topics is limited (both spatially and semantically) by the quality of the meta-analytic data in the automatically-extracted Neurosynth database (for discussion, see Yarkoni et al., 2011). In theory, greater specificity might be achievable using human-curated meta-analytic databases (e.g., BrainMap; Laird et al., 2005) or publicly deposited whole-brain images (Gorgolewski et al., 2015; Salimi-Khorshidi, Smith, Keltner, Wager, & Nichols, 2009). However, such resources are currently much smaller than Neurosynth--implying a significant decrement to the sensitivity of our data-intensive modeling approach--and, in the case of BrainMap, have usage restrictions that limit reproducibility and transparency. Nevertheless, it is clear that the present topics already converge closely with prior literature. Moreover, the integration of our topics with the public NeuroVault repository ensures that researchers will always be able to apply the most current topic sets to their data at the push of a button.

Second, the output of the GC-LDA model is necessarily data- and context-dependent. While the topics produced by the model generally have parsimonious interpretations that accord well with previous findings, they should be treated as a useful, human-comprehensible approximation of the true nomological network of neurocognition, and not as a direct window into reality. For the sake of analytical tractability, our model assumes a one-to-one mapping between semantic representations and brain regions, whereas the underlying reality almost certainly involves enormously complex many-to-many mappings. Similarly, re-running the GC-LDA model on different input data, with a different number of topics, or with different analysis parameters would necessarily produce somewhat different results. Of course, this concern applies equally to other large-scale data-driven approaches. We highlight it here simply because we would not want researchers to reify the topics we introduce here as if they are uniquely “real”. In our view, the overriding evaluation metric for any novel parcellation or clustering technique is whether it is scientifically productive over the long term (cf. Russell A. Poldrack & Yarkoni, 2015). With that caveat in mind, we believe that the framework introduced here strikes an excellent balance between interpretability, flexibility, and ease of use, and provides an important complement to previous data-driven approaches.

Lastly, while our decoding framework is formally Bayesian, the outputs it generates cannot typically be interpreted as probabilities, because the input images researchers conventionally seek to decode are mostly real-valued t or z maps whose meaning can vary dramatically. While this restriction limits the utility of our framework, it is, at present, unavoidable. Providing meaningful absolute estimates of the likelihood of different cognitive processes given observed brain activity would require either (a) that researchers converge on a common standard for representing observed results within a probabilistic framework (e.g., reporting the probability of subjects displaying supra-threshold activation in every voxel), or (b) re-training the GC-LDA model and associated decoding framework on a very large corpus of whole-brain images comparable to those that researchers seek to decode, rather than on a coordinate-based meta-analytic database. Of these two alternatives, we view the latter as the more feasible and productive strategy. We thus believe that the best hope for truly open-ended, fully probabilistic brain decoding lies in the widespread communal adoption of whole-brain images repositories like NeuroVault.org. We are optimistic that in the relatively near future, we will be able to use the topic modeling and decoding methods introduced here to produce highly informative, context-sensitive predictions about the mental processes implied by arbitrary patterns of whole-brain activity.

## Materials and methods

### Datasets

All data used to train the GC-LDA topic model came from the Neurosynth database (Yarkoni et al., 2011; neurosynth.org). The database contains activation coordinates that were automatically extracted from 11,409 published fMRI studies, as well as associated semantic terms extracted from the corresponding article abstracts. Further details have been reported in previous studies (Chang et al., 2012; Pauli, O’Reilly, Yarkoni, & Wager, 2016; Russell A. Poldrack et al., 2012; Yarkoni et al., 2011).

For decoding analyses, we used whole-brain maps obtained from several sources, including: (1) *The 500-subject release of the Human Connectome Project (Van Essen et al., 2013)*. We focused on single-subject whole-brain beta maps from several functional tasks. In all cases, we used experimental contrasts predefined by the HCP research team and included in the “preprocessed” data release (i.e., we did not preprocess or alter the provided contrast images in any way). Studied contrasts included the comparison between faces and shapes in the Emotion task; between language and math conditions in the Language-Math condition; between social and non-social motion in the Social Cognition task. (2) *NeuroVault.org maps*. We downloaded two sets of maps from the NeuroVault whole-brain image repository: (i) a completely random set of 100 images (subject to the constraint that each image had to come from a different image collection, to maximize independence of images), and (ii) a random set of 300 NeuroVault images that had been previously manually annotated using the Cognitive Atlas ontology for a completely different purpose (Sochat et al., in preparation). (3) *BrainMap ICA and Yeo et al. author-topic “cognitive component” maps*. We obtained these two sets of maps--reported in Smith et al. (2009) and Yeo et al. (2014), respectively--via the web.

### Topic modeling

A high-level schematic of the model we employ is presented in Figure 2; the model is presented using graphical model plate notation representation in Figure 8. We begin with the Neurosynth dataset, which contains data extracted from 11,406 published fMRI articles. Each of the 11,406 document consists of (1) a set of unigrams and bigrams of words extracted from the publication’s abstract, describing what each experiment was about, and (2) the set of peak-activation coordinates that were reported in HTML tables within the paper (for data extraction details, see Yarkoni et al, 2011). The model learns a set of *T topics*, where each topic is associated with some spatial distribution (e.g., a 3-dimensional Gaussian distribution with parameters *μ_t_* and *σ_t_*), and a multinomial distribution *ϕ_t_* over all of the unique types of linguistic features (consisting of unigrams and bigrams) in the corpus. This model is a generative model, meaning that it describes a process that can generate approximations of the observed data (the linguistic features and activation coordinates) via a set of latent (unobserved) topics.

**Figure 8.**
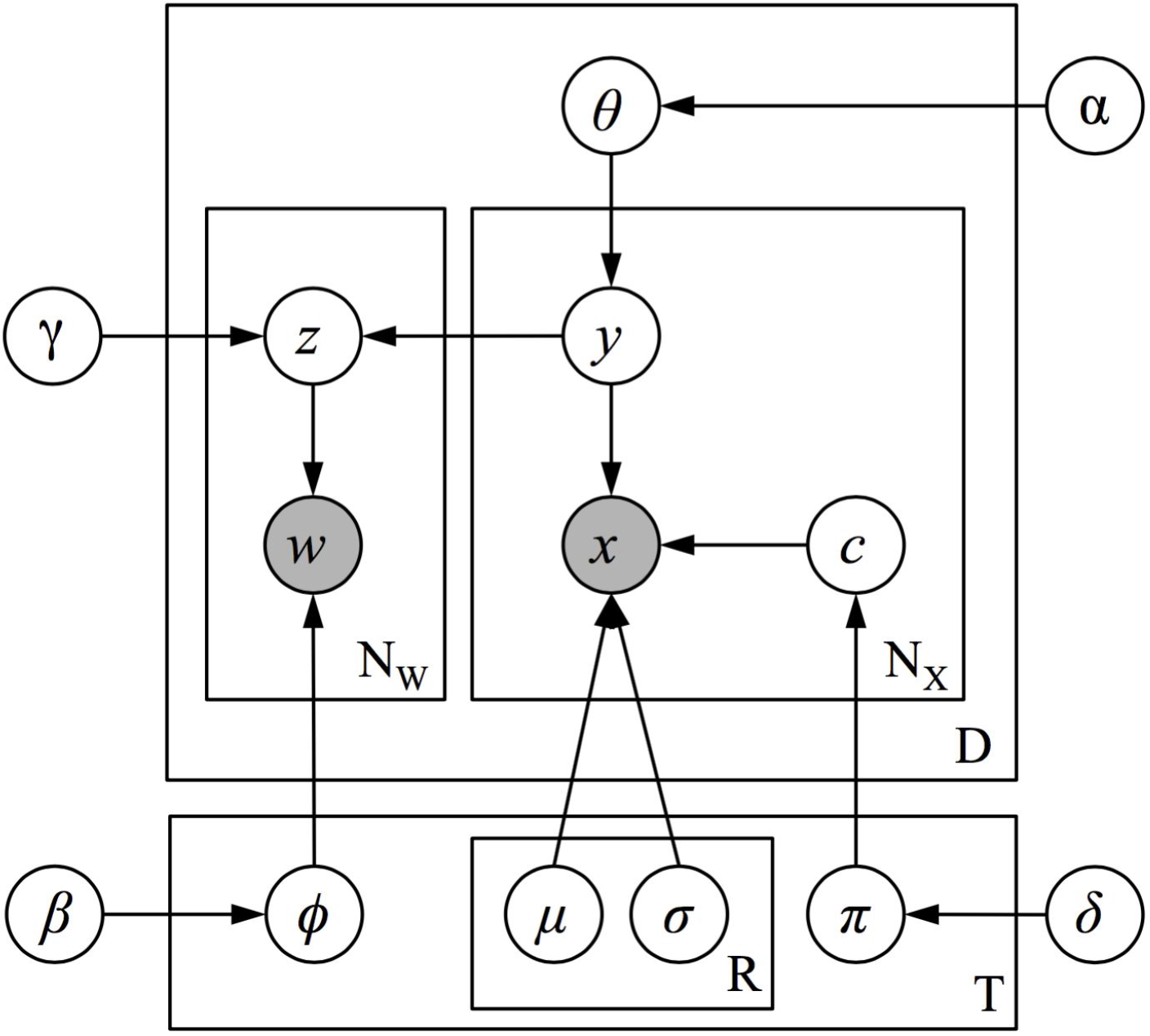
Graphical model of the full GC-LDA model.

The model assumes that each document *d* is generated by first sampling a multinomial probability distribution *θ_d_* over topics from a Dirichlet prior distribution. Then, to generate each activation peak *x* in the document, the document first samples a topic *y* from its distribution over topics *θ_d_* and then samples a peak activation at location *x* from the spatial distribution associated with topic *y*. To generate each word in the document, a topic *z* is sampled proportional to the number of times that the document sampled activations from each topic, and then a word token is then sampled from topic *z*’s probability distribution over word types *ϕ_z_*. To illustrate this process, consider the example Document 1 shown in Figure 2, which we can imagine describes an experiment measuring reaction times on a word-identification task. The model assumes that neural activation peaks reported in this experiment will be sequentially sampled from the spatial distributions associated with topics 1 and 2 (which relate to language processes and motor processes, respectively). The model then assumes that the words in the document--used to describe the experiment and its results--will be sampled from the linguistic distributions associated with topics 1 and 2, proportional to the number of times activation peaks were sampled from each of these topics.

Because the model enforces a correspondence between the frequency with which documents sample their words and activations from each topic, the model ensures that over the document corpus, the linguistic features associated with each topic will be closely related to the topic’s spatial distribution over activations. More specifically, the model will identify a topic-specific distribution over neural activations that tends to co-occur with the topic’s linguistic features across the corpus.

The general framework of the GC-LDA model allows the experimenter to choose any valid probability distribution for the spatial component of each topic. The results displayed in Figures 3 - 8 correspond to a GC-LDA model in which each topic’s spatial distribution is captured by a mixture of two Gaussian distributions that have been constrained to be symmetric about the x-axis. In our experiments, we evaluated several variations of the GC-LDA model using different probability distributions. We started with each topic having a single multidimensional Gaussian spatial distribution. We then replaced the single Gaussian distribution with a Gaussian Mixture distribution containing two components (i.e. subregions). In a further variant of this model (pictured in Figure 1), we constrained the spatial arrangement of the two component distributions of the Gaussian mixture distribution, such that their means were symmetrical with respect to the x-axis of the brain (i.e., so that for each topic, the spatial distribution would consist of one component region in the left hemisphere and a second component region in the right hemisphere). This allowed us to include an anatomical constraint based on known features of functional neuroanatomy—specifically the fact that there is generally a bilateral symmetry with respect to neural functionality. It further provided us with an automated way of measuring the lateral asymmetry of different cognitive functions (given by each topic’s probability of drawing an activation from its different components).

Given a formalized generative process for any of these models, we can use Bayesian inference methods to learn all of the latent (unobserved) parameters of this model from the observed data (see (Rubin, T. N., Koyejo, O., Jones, M. N., & Yarkoni, T., Submitted), for details). Specifically, the model learns a set of *T* topics, where each topic has an associated spatial probability distribution over the coordinates in the brain, as well as a multinomial distribution over linguistic features. The model additionally learns the topic mixture weights for each document.

### Text-to-image and image-to-text decoding

For text-to-image decoding (Fig. 7 A), we first compute a Topic (*T*) by word-type (*W*) matrix *P_W×T_* matrix of conditional probabilities, where cell *P_ij_* is the probability *P(t = i|w = j)* that the model assigns word type *w* from the text input to the *i^th^* topic. This matrix is computed, using Bayes’ rule, from the topic’s probability distributions over word types *Φ_W×T_* in the trained GC-LDA model, as follows:

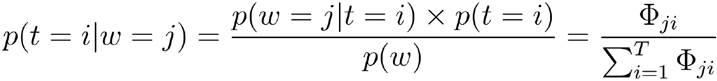

assuming a uniform prior probability of each topic, *p(t)*. We then obtain a vector of topic weights *τ* for the entire input by summing over the all word tokens *w* in the input; i.e.,

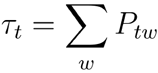

Lastly, we multiply this vector of topic weights by the topic x voxel matrix *A_T×V_* where cell *A_ij_* reflects the smoothed conditional probability *p(v = j|t = i)* that the model samples an activation at brain voxel *j* (of *V* total voxels) from topic *i*. The rows of this matrix are (smoothed versions of) the images displayed in Figure 3. The resulting (vectorized) whole-brain image is thus given by the product *τA*.

Note that the resulting values cannot be interpreted as probabilities, because we deliberately sum over words in the input rather than computing the joint probability. The reason for this is that, while the latter approach is technically feasible, and typically produces very similar results for short inputs, it produces unstable results when the input sentence exceeds a few words in length (because the sparse nature of the word-to-topic mapping results in the compounding of many very small probabilities).

For discrete coordinate-to-text decoding (Fig. 7B), we repeat the above process, but proceed in the opposite direction. That is, we first compute a Topic × Voxel matrix *P_T×V_*, where cell *A_ij_* reflects the conditional probability *p(t = i|v = j)* that the model assigns the the activation at voxel *j* in the input to the *i^th^* topic. This matrix is computed from the trained GC-LDA model by first computing the probability of sampling of each voxel from each topic (given each topic’s spatial distribution), and then renormalizing these probabilities using Bayes’ rule, under the same assumption used for text-to-image decoding of a uniform prior probability of each topic *p(t)*. We then sum over all of the input activations to obtain a vector of topic weights *τ_t_* for the given input. Lastly, we project the topic weights into the word space by multiplying the vector of topic weights *τ_t_* by the topic x word matrix *Φ_W×T_*, where cell *Φ_ij_*reflects the conditional probability *p(w = i|t = j)* that the model samples the *i^th^* word type from topic *j*.

To decode text from continuous whole-brain images (Fig. 7C), a slightly different approach is required. Although whole-brain decoding superficially resembles the decoding of discrete coordinates, the fact that the input images are real-valued and have arbitrary scaling precludes a true probabilistic treatment. Instead, we adopt a modified approach that weights the conditional probability matrix *P* by the similarity of the input image to each of the GC-LDA topic maps. We compute a vector of topic weights *τ* as:

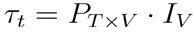

where *P_T×V_* is the topic × voxel matrix of conditional probabilities of assigning an activation at voxel *v* to topic *t*, and *I_v_* is the vectorized whole-brain input image. We then project the topic weights into word space in the same way as for the discrete coordinates (i.e., by computing *τ* · *Φ*). The scale of the resulting values is arbitrary, and depends on the input image, but the rank-ordering of terms is instructive and typically converges closely with human-annotated labels.

### Contextual decoding via topic “seeding”

Specifying priors on the GC-LDA topics can in principle be accomplished directly, by simply setting the desired prior probabilities on topics *P(t)* when computing the matrices *P_T×V_* and *P_W×T_*. The decoder results will then directly reflect the adjustment in both the text-to-image and image-to-text directions. However, researchers are unlikely to have strong intuitions about the relative base rates of the latent topics themselves. More commonly, they will instead wish to update the priors indirectly, based on a more intuitive expression of the experimental context or prior belief. This can be accomplished by “seeding” the priors with image and/or text inputs. In this case, the procedure can be thought of as a two-step application of the decoding methods described above. On the first pass, the input image or text is used to estimate values of *τ* (no further output is generated). On the second pass, the *τ* computed during the first pass is used as an informative prior *p(t)* in computing matrix *P_T×V_* or *P_W×T_* as described previously, and this updated matrix is applied to the actual image or text to be decoded. This procedure can repeat an indefinite number of times, as in a typical Bayesian context (i.e., the posterior *τ* probabilities become the priors for the next decoding application).

## Acknowledgments

The authors wish to thank Vanessa Sochat for contributing the manual NeuroVault annotations. This work was supported by National Institute Institute of Mental Health (NIMH) award R01MH096906 to TY, RAP, and MNJ.

## Supplementary Results

### Topic-based reconstruction of whole-brain maps

The probabilistic functional-anatomical atlas we introduce in the main text is useful not only for advancing theoretical understanding of the functional properties of different brain regions, but also for facilitating description and interpretation of novel whole-brain maps. One can conceptualize the topics produced by the GC-LDA model as a basis set that can be combined in various ways to produce much more complex patterns of whole-brain activity. In principle, a diverse array of whole-brain activation maps might be recaptured or reconstructed using a weighted combination of our regionally-circumscribed topics. This capability would provide a powerful means of reducing brain maps that nominally contain hundreds of thousands of distinct voxels to a much smaller number of meaningful, functionally distinctive units.

To test this idea empirically, we attempted to “reconstruct” a series of whole-brain activation maps by fitting a regression model that predicted activation at each voxel in each target image from activation levels in the 200 region-specific topic maps. We used a regression approach to reconstruct whole-brain activation maps using the topics generated by our GC-LDA model. First, we smoothed each of the 200 topic maps containing activation assignments (e.g., Figure 3b-c) with a 6 mm FWHM kernel. Next, we vectorized both the target whole-brain map and the 200 smoothed topic maps. We then fit an ordinary least squares regression model predicting the target whole-brain activation map from the 200 topic maps, i.e.,

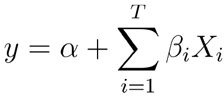

where *y* is the vectorized input image to decode, *α* is the intercept, *T* is the number of topics, *β_i_* is the estimated coefficient for the *i^th^* topic, and *X_i_* is the *i^th^* smoothed whole-brain topic map (cf. Figure 3). This analysis produces a set of 200 *β* coefficients that reflect the relative weight of each topic in the reconstructed/predicted map. We report the full model’s coefficient of determination (*R*^2^) as a metric of the model’s relative ability to describe the original map.

We applied this reconstruction approach to three very different sets of whole-brain images, including (1) the 20 BrainMap ICA maps reported by Smith et al. (2009); (2) a randomly-selected set of 100 maps drawn from the NeuroVault whole-brain image repository (Gorgolewski et al., 2015); and (3) single-subject activation maps from the the Human Connectome Project (HCP; Van Essen et al., 2013)---a landmark study that to date has released fMRI data from over 900 subjects performing a variety of experimental tasks (Barch et al., 2013). Supplementary Figure 4 illustrates reconstruction results for sample images of each type (for additional examples, see Supplementary Figures 5 - 7). For the BrainMap ICA images, reconstruction fidelity was almost universally high (mean R^2^ = 0.74), and visual inspection revealed striking similarity in the vast majority of cases (Supp. Fig. 4A, Supp. Fig. 5; the sole exception was component 19 [R^2^ = .23], which clearly consisted of artifactual activation on the fringe of the brain). For the NeuroVault maps---which varied widely in terms of task, analysis type, and sample size---reconstruction fidelity was somewhat lower (mean R^2^ = 0.46; Supp. Figure 4B), but the reconstructed maps preserved most of the spatial detail in the original maps (Supp. Fig. 7). As a general rule, maps sourced from clearly-defined group-level contrasts were easier to reconstruct than maps with ambiguous provenance.

In contrast, reconstruction accuracy was relatively poor for the single-subject HCP images, with mean R^2^ values ranging from 0.18 (social cognition task) to R^2^ = 0.3 (gambling task) across 4 different HCP tasks. This decrease was expected, however, as single-subject maps are necessarily noisier than group-averaged estimates, and also reflect considerable idiosyncracies in individual anatomy. Standard mass univariate group-level analyses are typically blind to such fine-grained differences, and can retain only the coarse patterns observed across the sample. The topic reconstruction approach can be viewed as an analogous means of regularizing low-level anatomical idiosyncracies and abstracting away high-level commonalities. That is, subjects whose neural responses to the same stimulus look very different in the original voxel space will typically have considerably more similar representations when reconstructed using our topics. For example, it is not at all apparent that the two single-subject images presented in Supp. Fig. 4C reflect the same functional task (i.e., the HCP Emotion task). By contrast, the topic-reconstructed images look much more similar, while still correlating strongly with each of the original two images.

### Supplementary Figures

[Note: Figures S1 and S7 are too large to include in this document, and are uploaded separately.]

**Figure S1** [uploaded separately]. Full results for all topics learned by the GC-LDA model. Each row represents a single topic. For each topic, the word cloud displays the top semantic associates (the size of each term is roughly proportional to the strength of its loading, and the orthviews display all hard assignments of activations to that topic (each point represents a single activation from a single study in Neurosynth).

**Figure S2.**
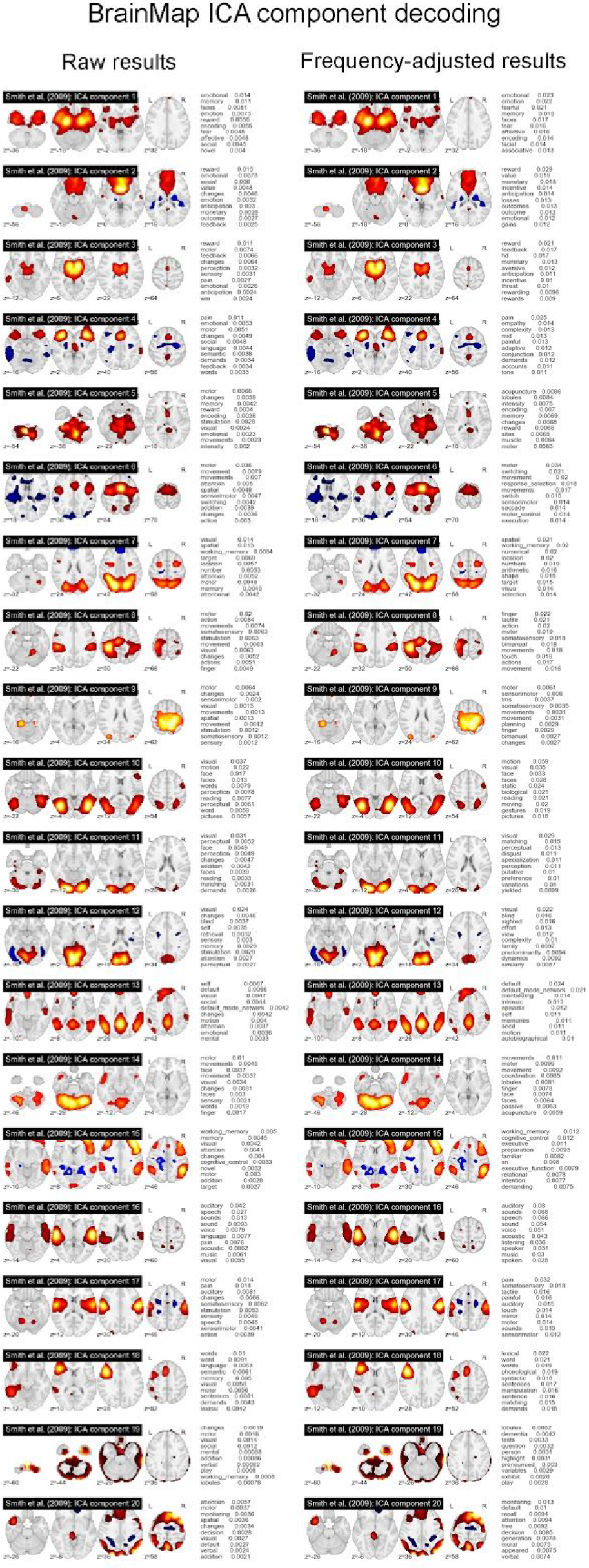
Topic-based decoding of 20 BrainMap-derived ICA components reported in Smith et al. (2009).

**Figure S3.**
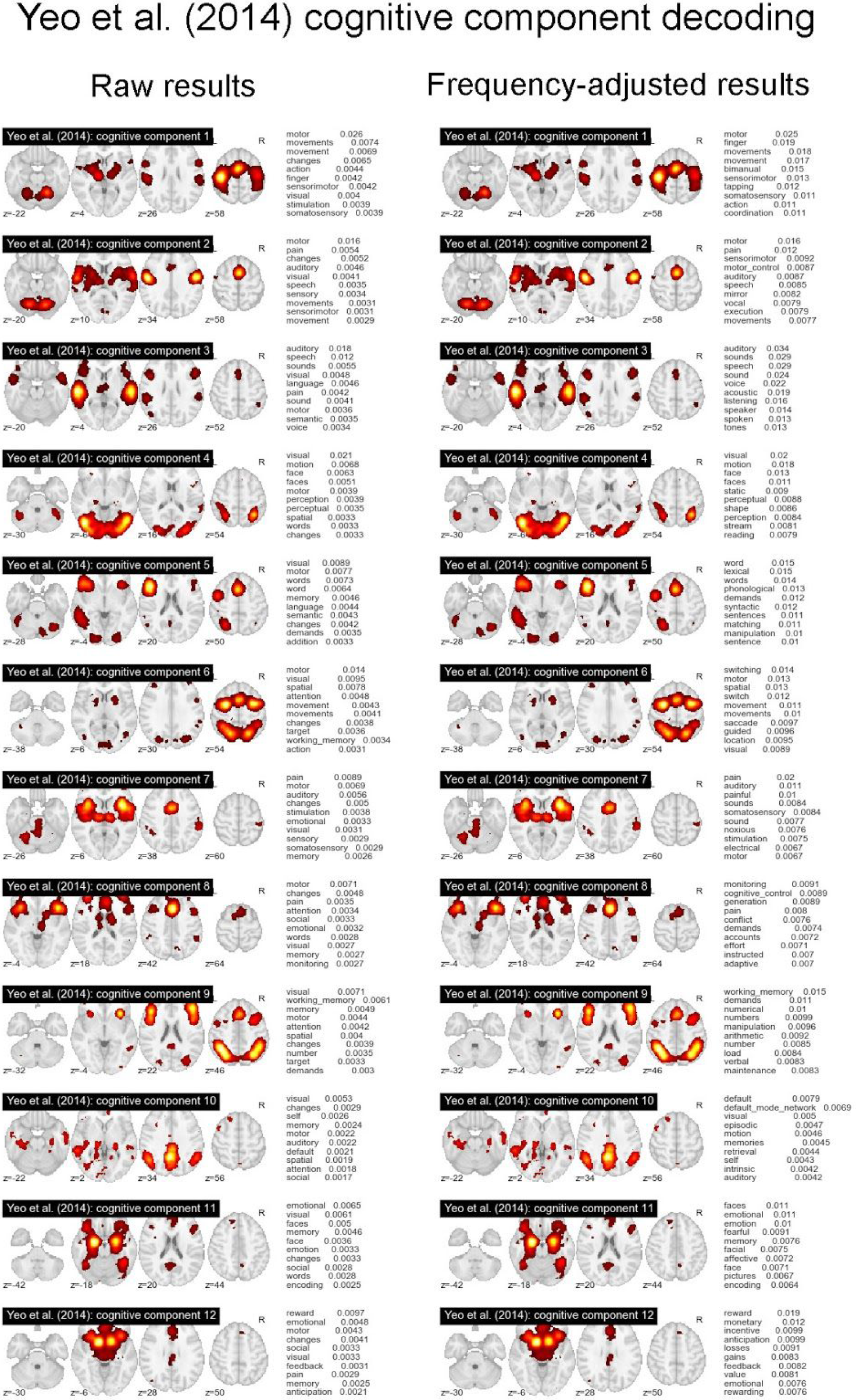
Topic-based decoding of 12 “cognitive components” reported in Yeo et al. (2014).

**Figure S4.**
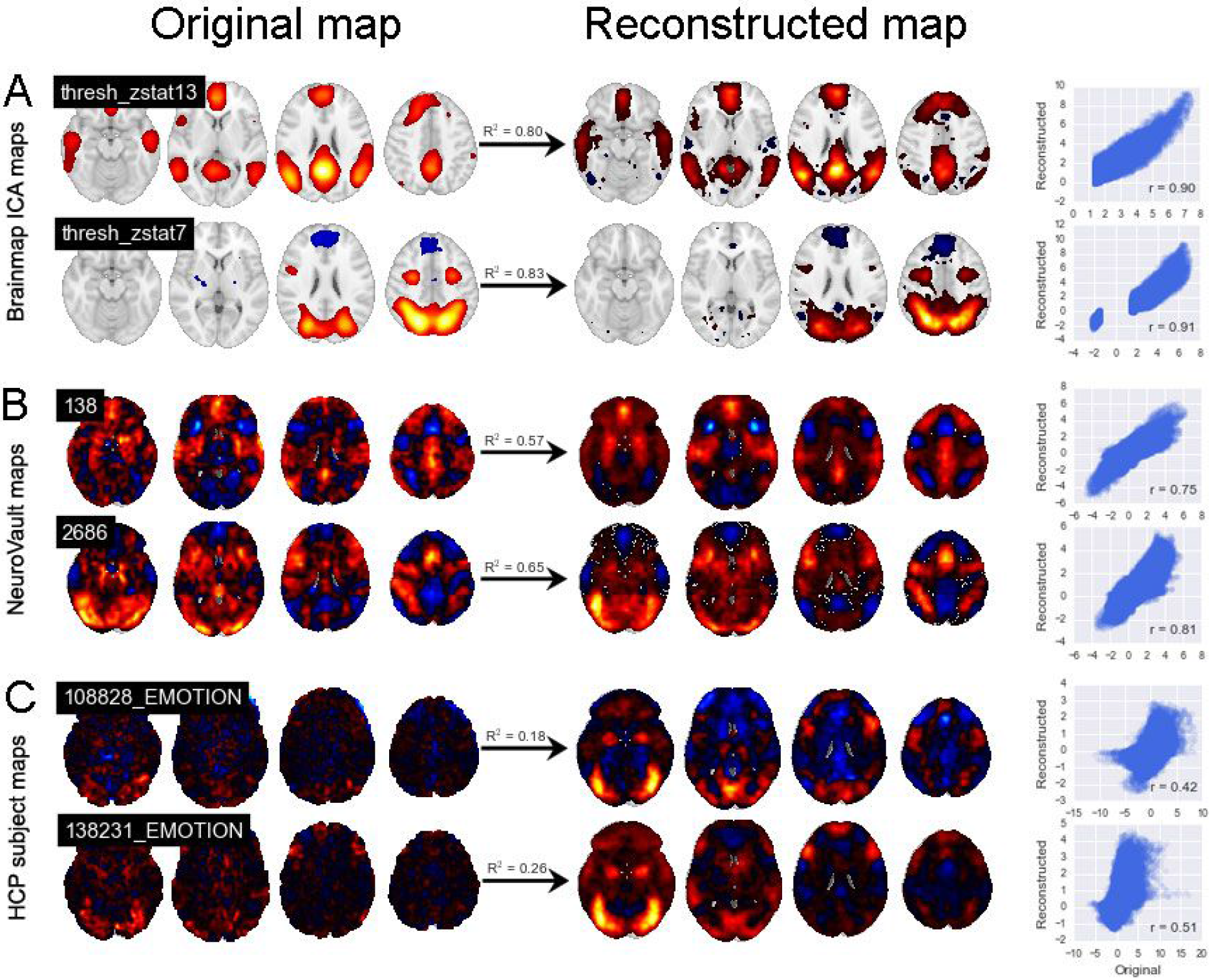
Topic-based reconstruction of whole-brain activity maps. Representative examples from (A) the set of 20 BrainMap ICA components reported in Smith et al. (2009), (B) the NeuroVault whole-brain image repository (Gorgolewski et al., 2015), and (C) single-subject contrast maps from the emotion processing task in the Human Connectome Project dataset (face vs. shape contrast). Each row displays the original (left) and reconstructed (center) image, along with the coefficient of determination (R^2^) for the fitted reconstruction model, and a scatter plot of all voxels (right).

**Figure S5.**
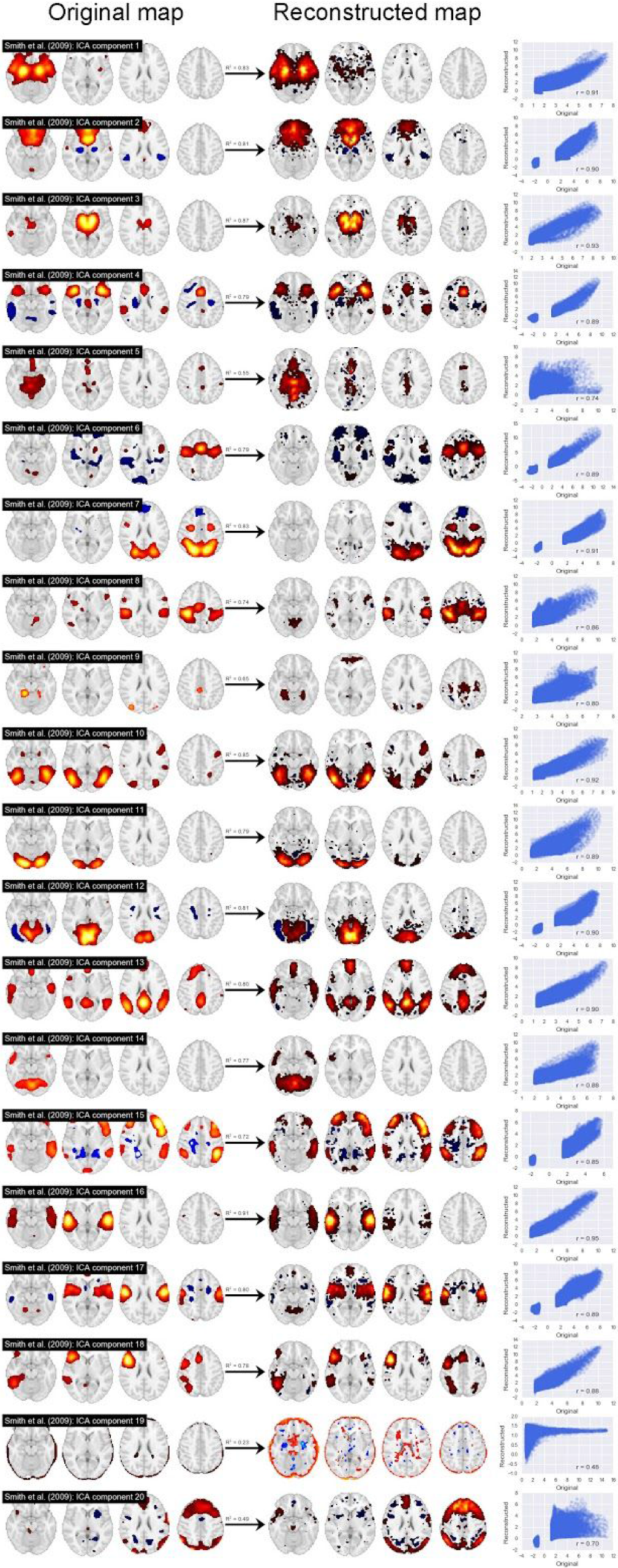
Reconstruction of 20 BrainMap ICA components reported in Smith et al. (2009).

**Figure S6.**
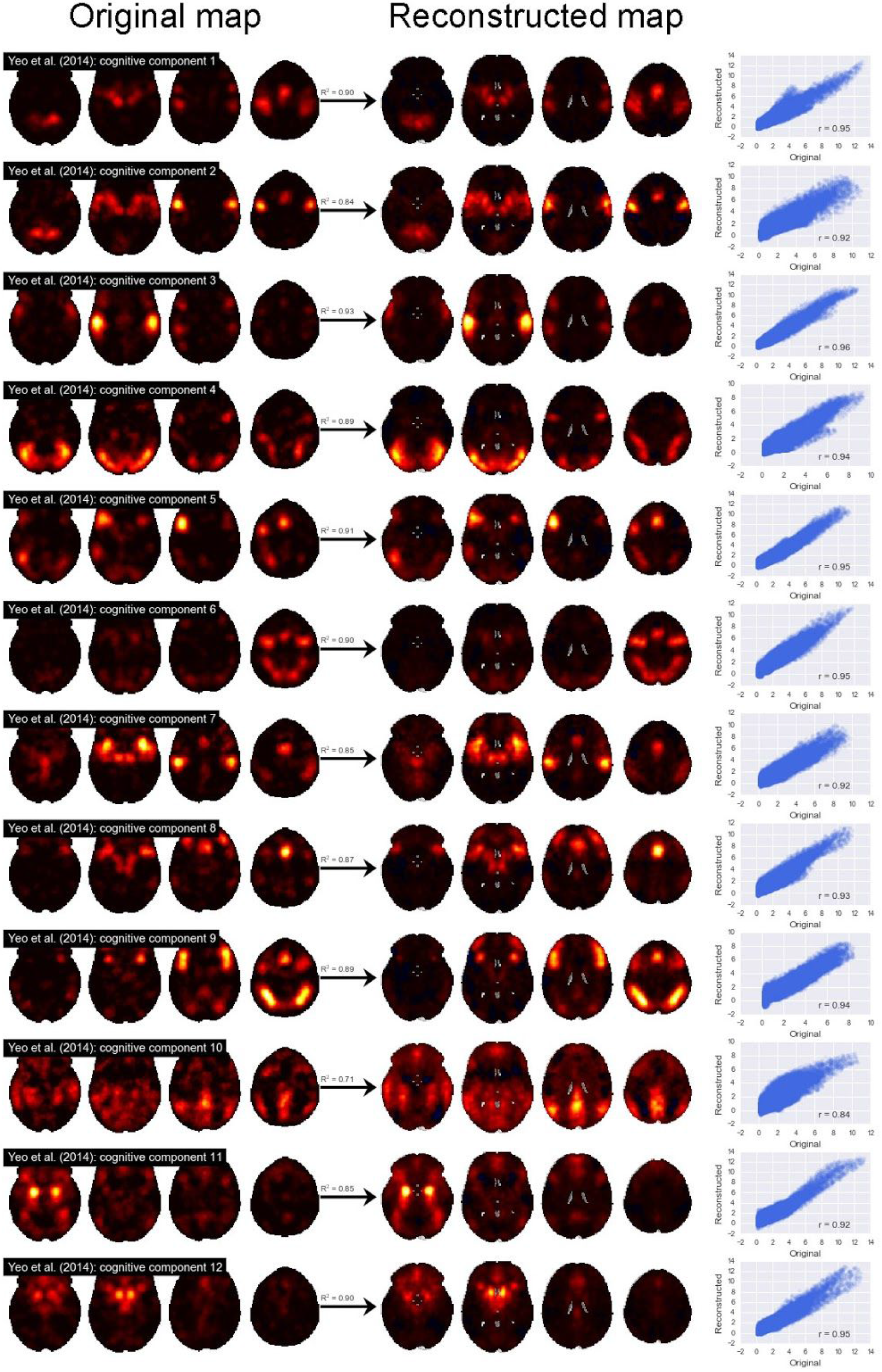
Topic reconstruction of 12 “cognitive components” reported in Yeo et al. (2014).

**Figure S7** [uploaded separately]. Topic reconstruction of 100 random maps extracted from the NeuroVault whole-brain image repository. Labels in white indicate human-annotated cognitive atlas paradigm, when available.

